# Illuminating the functional rare biosphere of the Greenland Ice Sheet’s Dark Zone

**DOI:** 10.1101/664334

**Authors:** Jarishma K. Gokul, Karen A. Cameron, Tristram D.L. Irvine-Fynn, Joseph M. Cook, Alun Hubbard, Marek Stibal, Matt Hegarty, Luis A.J. Mur, Arwyn Edwards

## Abstract

The Dark Zone of the western Greenland Ice Sheet is the most expansive region of contiguous bare terrestrial ice in the Northern Hemisphere. Microbial processes within the Dark Zone play an important role in driving extensive albedo reduction and amplified melting, yet the composition and function of those consortia have not been fully identified. Here we present the first results from joint 16S rRNA gene and 16S rRNA (cDNA) analysis for the comparison of input (snow), storage (cryoconite), and output (supraglacial stream water) habitats across the Dark Zone over the melt season. Our analysis reveals that all three Dark Zone communities are characterized by a preponderance of rare taxa exhibiting high protein synthesis potential (PSP). Furthermore, taxa with high PSP represent highly connected “bottlenecks” within community structure, consistent with roles as metabolic hubs within their communities. Finally, the detection of low abundance-high PSP taxa affiliated with *Methylobacterium* within snow and stream water indicates a potential role for *Methylobacterium* in the carbon cycle of Greenlandic snowpacks, and importantly, the export of potentially active methylotrophs to the bed of the Greenland Ice Sheet. By comparing the dynamics of bulk and potentially active microbial communities in the Dark Zone of the Greenland Ice Sheet our study provides insight into the mechanisms and impacts of the microbial colonization of this critical region of our melting planet.

## INTRODUCTION

Microbes that colonize snow and ice surfaces live at the critical interface between the atmosphere and cryosphere (Budyko 1969). Their potential to darken glacier surfaces and thereby amplify melt (Cook et al 2017, Lutz et al 2016, Nordenskiöld 1870, Ryan et al 2018, Stibal et al 2017, Takeuchi 2002), has been a long standing question that has recently been identified by the IPCC (AR5) as requiring urgent attention (IPCC 2014). The “Dark Zone” of the Greenland Ice Sheet is a conspicuous band of low albedo bare ice that covers some 10,000 km^2^ of the western ablating margin of the ice sheet. Surface melt rates of up to eight metres (water equivalent) per year have been observed here, representing a major component to the Greenland Ice Sheet’s negative mass-balance and contributor to global sea-level rise (Ryan et al 2016, Van Tricht et al 2016, Wientjes and Oerlemans 2010, Wientjes et al 2011). It is a biologically active surface (Cook et al 2012, Hodson et al 2010) where extensive microbial colonization drives regional surface albedo reduction and enhanced ablation (Ryan et al 2018, Stibal et al 2017, Tedstone et al 2017). Microbial processes associated with Greenland’s dark ice surface also contribute to the cycling and hydraulic export of microbial biomass (Cameron et al 2017, Dubnick et al 2017), organic carbon (Bhatia et al 2013a, Musilova et al 2017, Stibal et al 2010), nutrients (Bhatia et al 2013b, Hawkings et al 2016) in significant quantities to downstream englacial, subglacial, and proglacial hydrological networks and ecosystems which ultimately drain to the coast.

Previous studies of microbial diversity within the Dark Zone have focused on supraglacial communities within granular microbe-mineral aggregates termed cryoconite (Cameron et al 2015, Edwards et al 2014a, Musilova et al 2015, Stibal et al 2015) and glacier algae (Lutz et al 2018, Yallop et al 2012). These studies employed transects (e.g.(Edwards et al 2014b, Lutz et al 2018), or used pooled cryoconite material (Musilova et al 2015, Stibal et al 2015), thereby limiting detailed information regarding temporal bacterial community stability. Few studies have directly addressed the diversity of snowpack bacteria across the region (Cameron et al 2014). Despite the vast scale of this microbial habitat created by seasonal snowmelt (Ryan et al 2019), nothing is known of microbial temporal dynamics within this habitat. Furthermore, although fluvially-exported microbiota from the ice sheet surface may influence downstream biogeochemical processes such as subglacial methane cycling (Dieser et al 2014, Lamarche-Gagnon et al 2019), to our knowledge, the microbial diversity and functional potential of supraglacial meltwater exported from the Dark Zone remains undocumented.

The sequencing of 16S rRNA genes and 16S rRNA (reverse transcribed as cDNA) from co-extracted DNA and rRNA represents a common strategy within microbial ecology. Its application for the discrimination of “total” and “active” bacterial communities has been subject to critique, with limitations in the equivalence of rRNA and “activity” highlighted by Blazewicz et al (Blazewicz et al). With the caveat that ratios between 16S rRNA (cDNA) and 16S rRNA genes are indicative of protein synthesis potential (PSP; (Blazewicz et al 2013), rather than unequivocal quantitative evidence of contemporaneous growth, the technique offers the potential for insights into the responses of taxa to rapidly fluctuating environments. For example, within alpine proglacial streams which experience considerable diurnal fluctuation in temperature and discharge, joint 16S rRNA gene and 16S rRNA (cDNA) sequencing revealed that rare taxa were over-represented in the 16S rRNA (cDNA) population (Wilhelm et al 2014). Within the austere and isolated environs of the Dark Zone in summer, solar radiation, air temperature, melt intensity, and stream discharge all fluctuate with high periodicity, typically diurnally, and hence well within the typical doubling times of supraglacial microbes (Anesio et al 2010, Williamson et al 2018). How the bacterial communities of the Dark Zone respond to these fluctuations remains unknown, and the potential for these rare taxa to disproportionately influence community structure is unknown.

In this study, we address these questions by presenting an integrated study of community structure, connectivity, and its functional potential within three principal bacterial habitats within the Dark Zone: snow (input), cryoconite hole (storage) and runoff (output). We evaluate the temporal dynamics of bacterial communities from these three habitats using analysis of both 16S rRNA gene and 16S rRNA (cDNA). This was performed by sampling at weekly intervals, in June and July 2014, to incorporate the transition from early season melt onset to snow-free exposed ice surface.

## MATERIALS AND METHODS

### Methods summary

Sampling took place on the western ablating margin of the Greenland Ice sheet, within the Dark Zone and adjacent to the Kangerlussuaq (K-) transect S6 automatic weather station (AWS), at 67°05’N, 49°23’W; 1020 m asl (FIGURE 1). Cryoconite, snow and water from supraglacial meltwater streams was collected in triplicate, on seven sampling occasions, at weekly intervals between June 19 and July 31 2014. Independent sites, within a 25m^2^ area were used for each sampling occasion. A total of 62 samples were collected and chemically preserved using Soil Lifeguard (MO BIO Laboratories). Samples were then frozen as soon as possible. Upon return to the home laboratory, community DNA and RNA was co-extracted from snow and meltwater samples previously concentrated on 0.22 µm Sterivex GP polyethersulfone filters (Millipore, MA, USA) using a modified PowerWater® Sterivex™ DNA Kit and from cryoconite using a PowerBiofilm^™^ RNA Isolation Kit (MO BIO Laboratories) prior to 16S rRNA gene and 16S rRNA (cDNA) quantitative PCR (qPCR) and V3-V4 region MiSeq (Illumina) sequencing. All sequence data are available on EBI-SRA under the study accession number PRJNA318626. The methods employed for sample archival, nucleic acid extraction, qPCR, sequencing and data processing are detailed in full as supplementary methods.

**FIGURE 1:**
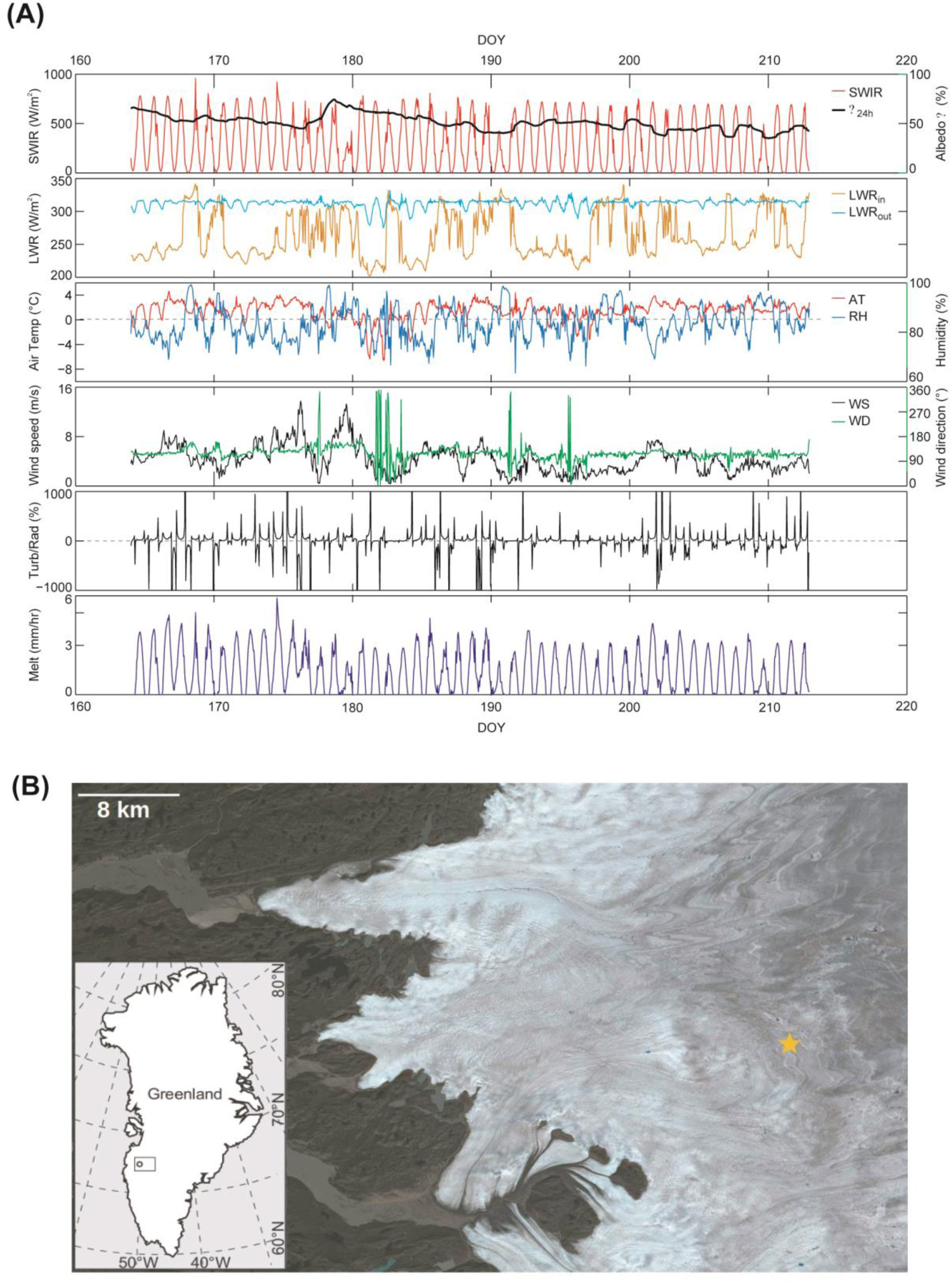
Overview of the study location and physical conditions. (A) The Dark Zone of the Greenland Ice Sheet. RGB composite image of the Kangerlussuaq region of the Greenland Ice Sheet generated from the European Space Agency Sentinel 2 reflectance product (atmospheric correction and reprojection of Level 1C tile downloaded from earthexplorer.usgs.gov using Sen2Cor, then bands 2, 3 and 4 merged and scaled using GDAL) showing the study site marked with a star, 38 km inland of the Greenland Ice Sheet margin. (B) Shortwave Incident Radiation (SWIR), Long Wave Radiation (LWR), air temperature, humidity, wind speed and direction, turbulent radiation (Turb Rad) and melt rate monitored at automatic weather station site S6. Meteorological data courtesy of CJPP Smeets and MR van den Broeke, Utrecht University.

## RESULTS

### Bacterial 16S rRNA gene and 16S rRNA (cDNA) quantification

Quantitative PCR was used to analyse the amplifiable copy number of 16S rRNA genes and 16S rRNA in DNA and cDNA samples (Figure 2). The abundance of the cryoconite bacterial community appeared highly consistent across the sampling period (Figure 2A) with weekly averages of 2.4 - 4.5 × 10^5^ amplifiable copies of the 16S rRNA gene per gram dry weight of cryoconite. The amplifiable copy number of the 16S rRNA pool fluctuated, ranging between a weekly average of 2.3 × 10^7^ (week 4) and 1.6 × 10^9^ (week 2) per gram dry weight of cryoconite. The ratio between 16S rRNA gene and 16S rRNA (cDNA) amplifiable copy number is interpreted as a marker of the overall bacterial PSP. Average cryoconite bacterial PSP showed high (1:88, week 4) to extremely high ratios (1:5000, week 2) throughout the study period. This is indicative of a high level of potential activity relative to biomass within the cryoconite bacterial communities sampled.

**FIGURE 2:**
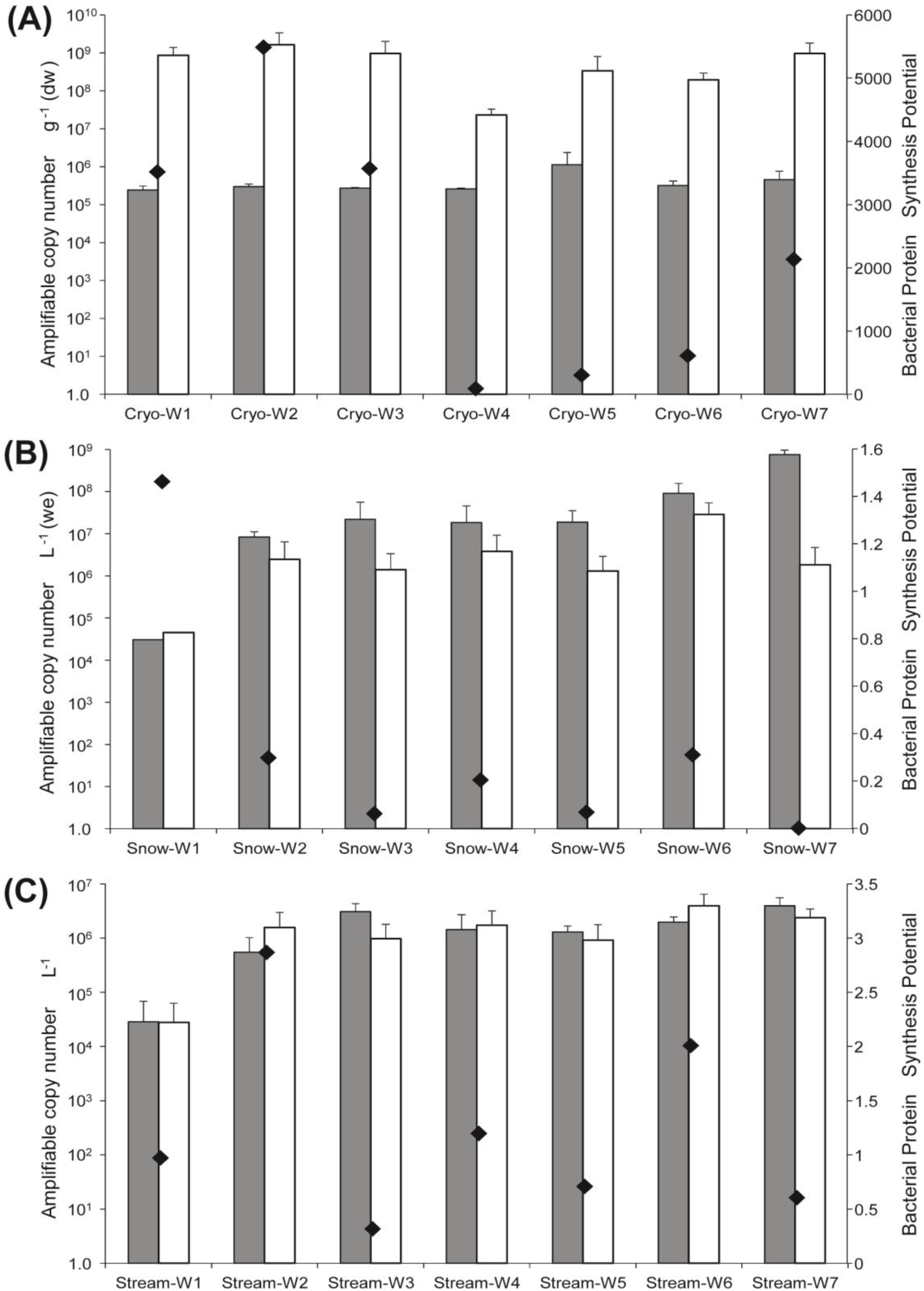
Quantitative PCR data on 16S rRNA gene (grey bars) and 16S rRNA (cDNA; open bars) amplifiable copy number and bacterial protein synthesis potential (diamonds) for (A) cryoconite, (B) snow, and (C) meltwater based upon analysis of triplicate weekly samples for each habitat for 7 weeks after the 19^th^ of June 2014. Note the different scales for each sub-panel.

For the snow bacterial community (Figure 2B), weekly average amplifiable copies of the 16S rRNA gene increased from 3.0 × 10^4^ copies per litre (water equivalent) in the first week of the study to 7.6 × 10^8^ copies per litre (water equivalent) in the final week. Weekly average amplifiable copies of 16S rRNA (cDNA) also increased from 4.5 × 10^4^ copies per litre (water equivalent) in the first week of the study to 2.8 × 10^7^ copies per litre (water equivalent) in the penultimate week. Weekly average bacterial PSP values for snow were consistently below equivalence, with exception of the first week (1:1.5 ratio). When viewed in the context of the rapid seasonal wastage of the snowpack in the study period, this likely represents the melt scavenging of bacterial cells incurring physical accumulation of quiescent biomass within the residual snow pack.

Stream water bacterial communities (Figure 2C) exhibited 16S rRNA gene amplifiable copy numbers several orders of magnitude lower in week 1 (2.8 × 10^4^ 16S rRNA gene copies per litre) compared to the remainder of the study period (3.1 - 3.6 × 10^6^ 16S rRNA gene copies per litre). rRNA copy numbers were at least an order of magnitude higher (5.5 × 10^5^ to 3.9 × 10^5^ 16S rRNA copies per litre) compared to the first week. Stream water bacterial PSP values varied between 0.3 and 2.8 during the study period. In summary, it is likely the stream water bacterial community was an admixture of quiescent and active taxa in transit from different sources (e.g. snowmelt, ice melt, cryoconite and other biofilms) from the ice sheet surface.

### Community structure

The total number of reads obtained after sequence processing was 2,673,556 with a maximum of 130,001 reads (GrIScDNAcryo6.3), a minimum of 5 (GrISstream3.1) and a mean of 21,561 reads. Sequences were filtered and rarefied to 943 sequences per sample, resulting in the exclusion of 1 cryoconite DNA sample, 2 stream DNA samples, 3 cryoconite cDNA samples, 3 snow cDNA samples and 6 stream cDNA samples from downstream analysis. Sequences were clustered into 566 operational taxonomic units (OTUs) at 97% sequence similarity. 13.59 % of 16S rRNA gene OTUs and 14.40 % 16S rRNA (cDNA) OTUs were common to all three habitats.

Over 99 % of the sequences in the dataset from cryoconite hole, snow and stream water habitats were successfully assigned to the Greengenes taxonomy using UCLUST. .Non-metric multidimensional scaling of OTUs clearly ordinated both 16S rRNA gene and 16S rRNA (cDNA) profiles of the bacterial communities by habitat type (Figure 3A). These trends were confirmed by PERMANOVA of fourth-root transforms of Bray-Curtis distances of OTU relative abundance matrices. Highly significant differences were found between all habitat types (Pseudo*F =* 13.8, *p =* 0.0001, Supplementary Table 2). Furthermore, highly significant differences are apparent between OTU composition of 16S rRNA gene and 16S rRNA (cDNA) profiles for each of the habitat types (Pseudo*F* values *=* 5.3 - 22.5, *p =* 0.0001). The snow and stream communities revealed in 16S rRNA gene profiles (Pseudo*F =* 2.6, *p =* 0.0001 and Pseudo*F =* 2.4, *p =* 0.0001 respectively) were temporally dynamic at weekly sampling resolution, with highly significant differences. The 16S rRNA (cDNA) profiles of cryoconite and snow were significant and highly significantly different by week (Pseudo*F =* 1.7, *p =* 0.02 and Pseudo*F =* 3.0, *p =* 0.0004 respectively), while stream profiles are temporally stable. In contrast, while cryoconite exhibited highly significantly different 16S rRNA gene and 16S rRNA (cDNA) profiles, the 16S rRNA gene profiles were temporally stable. All PERMANOVA results are detailed in Supplementary Table 2.

**FIGURE 3:**
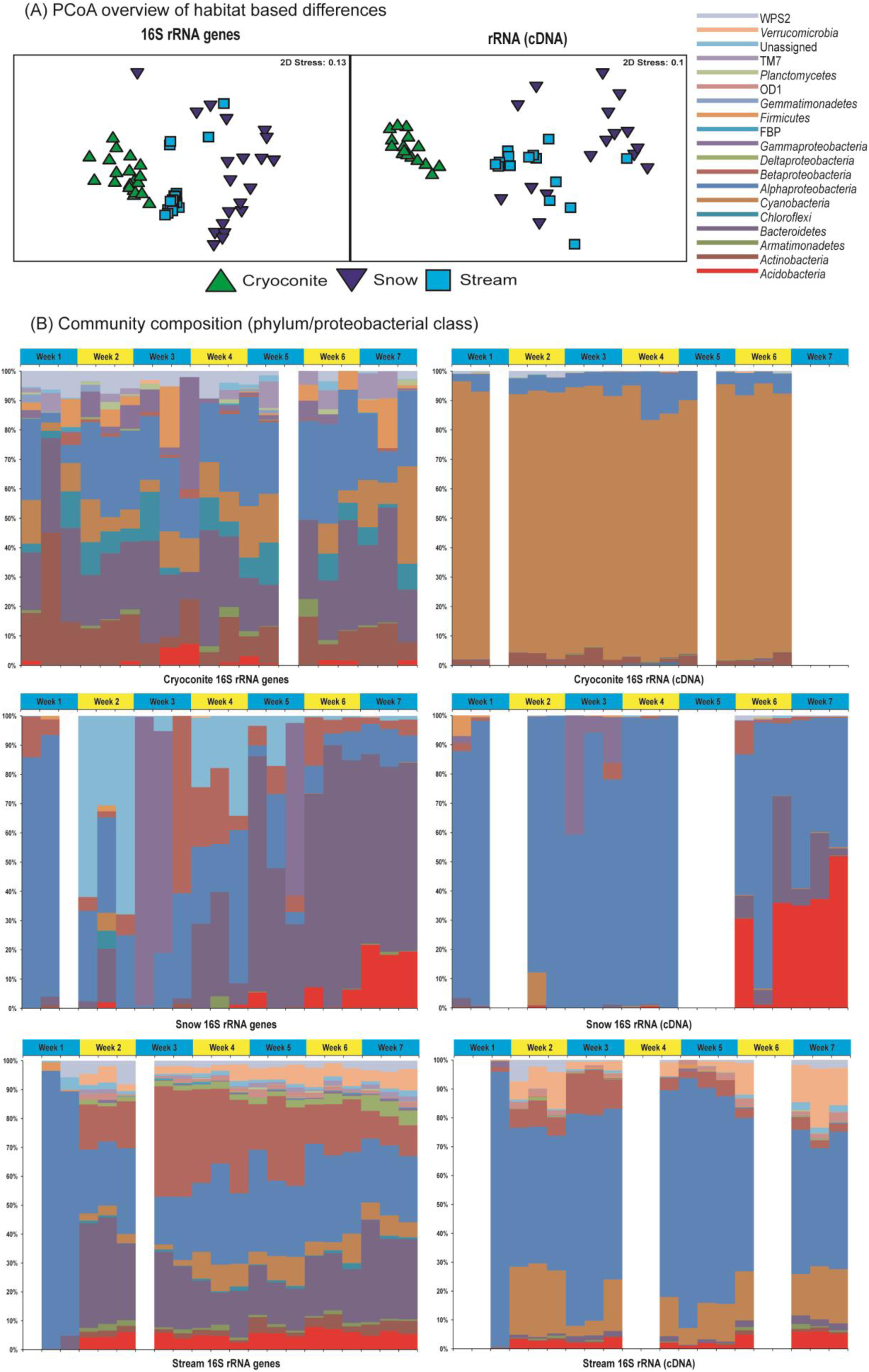
Overview of amplicon sequencing data. (A) Non-metric multi-dimensional scaling (nMDS) of fourth-root transformed Bray Curtis distances in OTU relative abundances for 16S rRNA genes and 16S rRNA ordinated by habitat. (B) Community composition based on phylum (or proteobacterial class) relative abundance for 16S rRNA genes and 16S rRNA ordinated by habitat. Blank bars indicate samples excluded upon rarefaction.

### Trends in taxonomic composition

Pronounced differences between 16S rRNA gene and 16S rRNA (cDNA) taxonomic profiles are apparent for each of the habitats (Figure 3B). Whereas 16S rRNA gene data reveal Actinobacteria, Bacteroidetes and Alphaproteobacteria are the major groups in cryoconite with a modest representation from Cyanobacteria, from the 16S rRNA (cDNA) data Cyanobacteria are the strikingly dominant group throughout the study period (Figure 3). Similarly, while the 16S rRNA gene profiles of snow reveal a transition between Alphaproteobacteria dominated community to a Bacteroidetes dominated community during the study period, Alphaproteobacteria remains the dominant group within the 16S rRNA (cDNA) profile, with an increase in Acidobacteria in the final two weeks of the study. Meanwhile, the discordance between 16S rRNA gene and 16S rRNA (cDNA) profiles are further mirrored in supraglacial streamwater, where Bacteroidetes and Betaproteobacteria were found to be in equitable dominance in the 16S rRNA gene dataset, and Alphaproteobacteria strongly dominated the 16S rRNA (cDNA) profiles. In summary, the trends in taxonomic composition observed are consistent with discrete bulk and potentially active communities, with phototrophic cyanobacteria active relative to biomass in cryoconite, and the Alphaproteobacteria notably active in snow and stream water.

We considered the potential impact of contamination on the taxa detected in our samples. Negative controls comprising blank DNA extractions were sequenced in parallel with field samples. The controls returned a small number of reads assigned to a total of 10 taxa, and were therefore excluded from the analysis of rarefied data. The most abundant sequence in the control samples was a *Salinibacter* sp. represented by a total of 16 reads (Supplementary Table 3). In contrast, the thirty most represented OTUs from 16S rRNA gene and 16S rRNA (cDNA) data from each habitat match taxa detected orthogonally from the natural environment, with glacial and other cryospheric habitats strongly represented (Supplementary Table 4). The influence of contamination on the present study is therefore considered minimal.

### 16S rRNA gene sequencing

#### Trends in potential activity and relative abundance

The strikingly distinctive 16S rRNA gene and 16S rRNA (cDNA) profiles were further investigated to reveal taxonomic groups over- and under-represented in 16S rRNA (cDNA) compared to 16S rRNA genes (Figure 4). All detected phyla/proteobacterial classes with the exception of *Cyanobacteria* were under-represented in 16S rRNA (cDNA) for cryoconite (Figure 4A), whereas Alphaproteobacteria, Acidobacteria, Firmicutes and WPS2 were over-represented in snow (Figure 4B) and Alphaproteobacteria, Verrucomicrobia, OD1 and Firmicutes were over-represented in stream water communities (Figure 4C).

**FIGURE 4:**
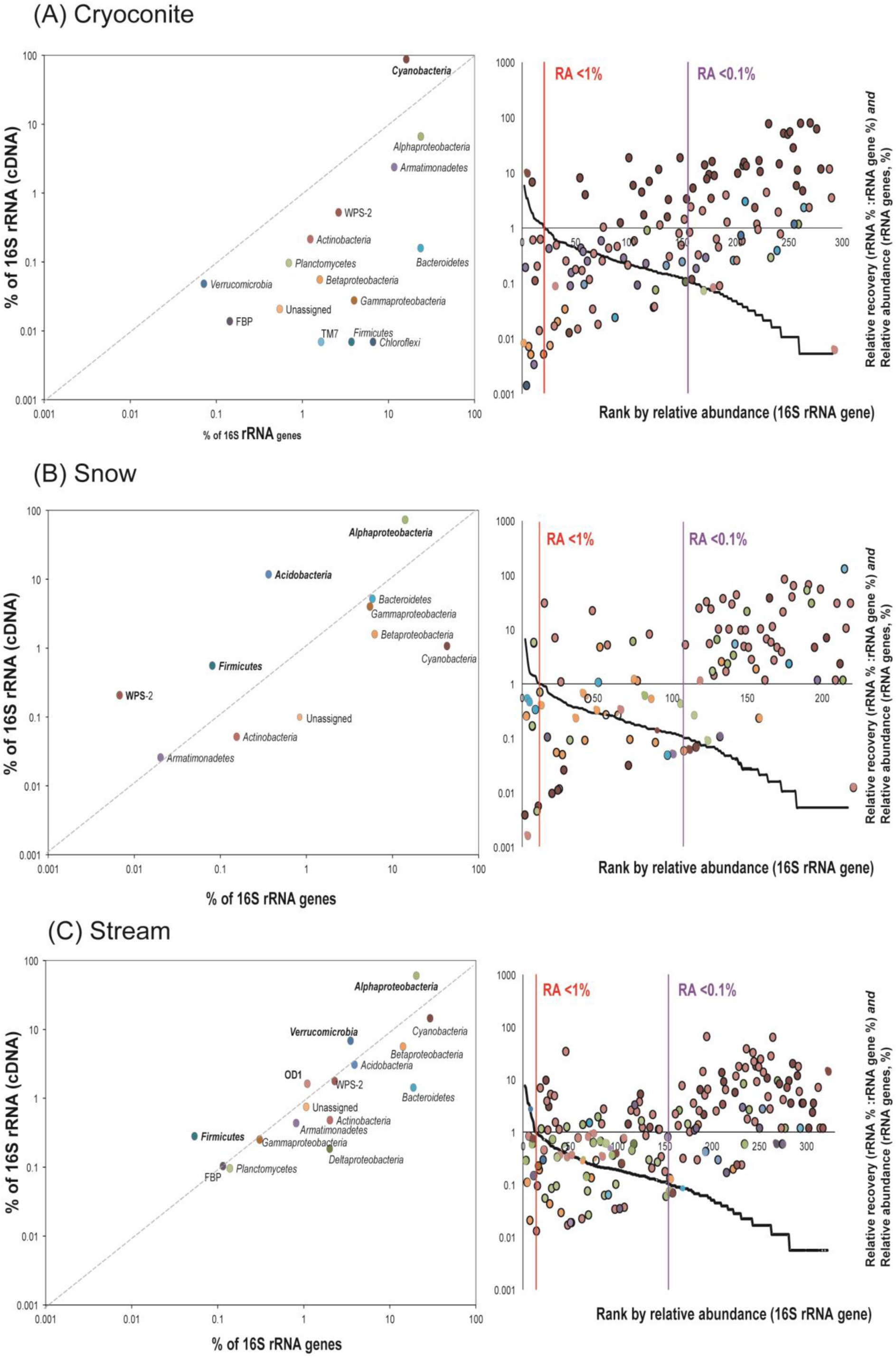
Relationships between 16S rRNA gene and 16S rRNA (cDNA) data for (A) cryoconite, (B) snow, and (C) streams during the study period. For each habitat, the correlation between phylum (and proteobacterial class) relative abundances between 16S rRNA gene and 16S rRNA (cDNA) data is shown. The diagonal line is used to indicate 1:1 equivalence. Rank abundance curves denote the OTU level distribution of taxa. The black line indicates the rank abundance curve of 16S rRNA gene OTUs against their relative abundance (left vertical axis). Data points represent individual OTUs, coloured by their parent phylum or proteobacterial class. Vertical lines indicate the position of taxa below 1% (red) and 0.1% (purple) relative abundance (RA).

OTUs present at ≤ 1% relative abundance constituted a large percentage of the bulk community within rank abundance curves (Fig 4A-C). Applying the ≤0.1 % of relative threshold commonly used to delimit the “rare” biosphere (Pedrós-Alió 2012), 57 % of cryoconite OTUs, 62 % of snow OTUs and 63 % of stream OTUs would be considered “rare” taxa within the Dark Zone bulk community. Rare taxa would need to exhibit a minimum mean relative abundance of ≥0.005% to be represented in datasets of this size. Rank abundance curves show that 48 % of rare cryoconite-habitat OTUs, 40.45 % of rare snow-habitat OTUs and 42.36 % of rare stream-habitat OTUs exhibit positive protein synthesis potentials (PSP, the ratio between 16S rRNA gene and 16S rRNA [cDNA] relative abundance) over the course of 7 weeks. In each community (Figure 4 A-C), taxon PSP is negatively correlated with mean taxon relative abundance (Spearman correlation; cryoconite: *r =* −0.63, *p <* 0.0001, snow: *r =* −0.65, *p <* 0.0001, stream *r =* −0.55, *p <* 0.0001). The trends exhibited are congruent with the notion that certain rare taxa in surface habitats in Greenland’s Dark Zone exhibit disproportionately high protein synthesis potential.

#### Dynamics of high-PSP OTUs

To establish the contribution of high PSP OTUs over time, OTUs exhibiting weekly PSP averages ≥1 in the dataset were plotted over time and compared to their relative abundance within the 16S rRNA gene dataset (Figure 5). For the cryoconite community (Figure 5A), 30 of 34 OTUs meeting this criterion were members of Cyanobacteria, with taxa assigned to *Leptolyngbya* representing the majority, including both the highest PSP OTU (*Leptolyngbya-*76) and highest relative abundance OTU (*Leptolyngbya-*3). In the snow community, Alphaproteobacteria represented 22 of 30 taxa with weekly average PSP ≥ 1 (Figure 5A). Notably, *Methylobacterium-*1 is highly abundant in the first week of the study with a corresponding mean PSP of 6.8. However, for the next three weeks, while *Methylobacterium*-1 shows much lower relative abundance, its PSP is strikingly high (ranging 185- to 304-fold).

**FIGURE 5:**
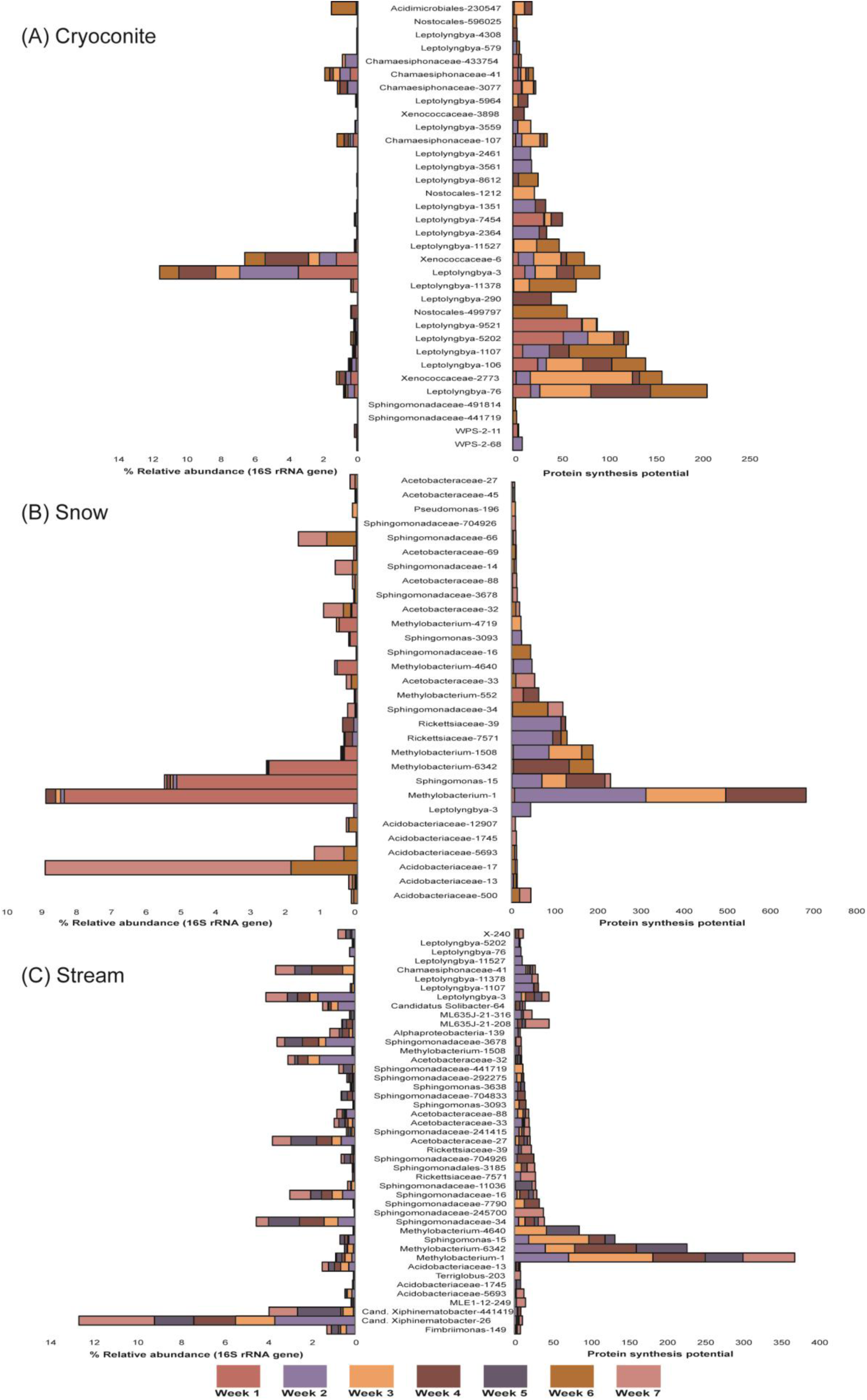
Temporal dynamics of OTUs with average weekly Protein Synthesis Potential ≥1 for (A) cryoconite, (B) snow, and (C) streams during the study period. The left hand plots show the relative abundance of each taxon by week while the right hand plots show the protein synthesis potential. Each OTU is named according to the lowest grade Greengenes taxon assigned and its denovo-OTU reference.

*Methylobacterium*-1 is not detected in snow community after week four. In all, six OTUs assigned to *Methylobacterium* are prominent in the high PSP taxa of the snow community. This is echoed within the stream community. Here, *Methylobacterium*-1 again shows high PSP values, in the range of 49 - 111 between weeks two and seven, however its relative abundance is low, amounting to <2 % of the community overall. Four OTUs assigned to *Methylobacterium* are present among 28 Alphaproteobacteria OTUs, with Sphingomonadaceae taxa well represented. In all, 45 OTUs show weekly average PSP ≥1. Seven *Cyanobacteria* affiliated with *Leptolyngbya* (including the *Leptolyngbya-*3 and *Leptolyngbya*-76 prominent in the cryoconite community) are present with four members of Acidobacteria. The prominence of high PSP taxa in stream water from lineages conspicuous within the snowpack and cryoconite community is consistent with the runoff export of potentially active taxa from surface habitats of the Greenland Ice Sheet’s Dark Zone.

#### Keystone species-high PSP rare OTU relationship

Taxa that exert a disproportionate influence on the structure of the microbial community, despite low or moderate abundances can be termed keystone species (Power and Mills 1995). High betweenness centrality, measured as the shortest number of paths between any two other OTUs passing through that OTU, is interpreted as a hallmark of a keystone species (Peura et al 2015). Co-occurrence analysis identified sixteen OTUs with betweenness-centrality scores (in the range 7.16 to 0.2; Table 1). All have cumulative positive PSP ratios in at least one habitat over the course of the study, and with the exception of one *Comamonadaceae* OTU with a cumulative mean PSP of 1.98, the remainder show high to very high maxima in their cumulative PSP ratios, in the range 10.5 - 405.8. Again, *Methylobacterium-*1 is represented, with the highest (snow: 405.8) and second highest (stream: 368.2) cumulative mean PSPs. Two other OTUs assigned to *Methylobacterium* show the next highest cumulative mean PSPs in stream and snow. For cryoconite, *Leptolyngbya* assigned OTUs are prevalent as keystone taxa with high cumulative mean PSP. Of the four *Leptolyngbya* assigned OTUs, *Leptolyngbya*-76, *Leptolyngbya*-106 and *Leptolyngbya*-3 show the highest cryoconite PSPs, but have modest betweenness scores. The considerable overlap between putative keystone species and high PSP taxa, including those present at low abundance, presents the possibility that taxa with high levels of protein synthesis potential are influential in the dynamics of their communities irrespective of their relative abundance. Thirteen of the sixteen OTUs are most closely related to taxa distributed across the global cryosphere (Table 1).

**Table 1:** Cumulative Protein Synthesis Potential (PSP) and relatives of keystone OTUs present in 16S rRNA gene profiles of cryoconite (Cryo), snow and stream habitats

|  |  |  |  |  |  |  |  |  |  |  |
| --- | --- | --- | --- | --- | --- | --- | --- | --- | --- | --- |
| Greengenes-OTU | Phylum | Betweenness | Cryo-PSP | Snow-PSP | Stream-PSP | Relative | Taxon | Accession | % ID | Habitat |
| *Sphingomonadaceae* -14 | Proteobacteria | 7.17 | 1.59 | **10.54** | 3.93 | CER | Uncultured bacterium clone Bysf-47-Sf10-014 | HQ622730.1 | 99 | Austre Lovenbreen, Svalbard - glacier ice core |
|  |  |  |  |  |  | CNR | *Sphingopyxis* sp. JJ2203 | JX304649.1 | 98 | Jangsu-gun, South Korea - lake |
| *Comamonadaceae* -5 | Proteobacteria | 6.25 | 0.25 | **1.89** | 1.30 | CER | *Polaromonas* sp. MDB2-14 | JX949585.1 | 99 | China - glacier |
|  |  |  |  |  |  | CNR | *Polaromonas* sp. MDB2-14 | JX949585.1 | 99 | China - glacier |
| *Methylobacterium* -1 | Proteobacteria | 4.99 | - | **405.88** | 368.26 | CER | Marine bacterium MSC10 | EU753147.1 | 99 | Nahant, Massachusetts - tidal flat sand |
|  |  |  |  |  |  | CNR | Marine bacterium MSC10 | EU753147.1 | 99 | Nahant, Massachusetts - tidal flat sand |
| *Sphingomonadaceae* -34 | Proteobacteria | 4.75 | 4.63 | **118.83** | 39.16 | CER | Uncultured bacterium clone QA4_1_042 | LC076727.1 | 99 | Qaanaaq Glacier, Greenland - cryoconite granules |
|  |  |  |  |  |  | CNR | *Novosphingobium* sp. STM-24 | LN890294.1 | 96 | Freshwater |
| *Acetobacteraceae* -27 | Proteobacteria | 2.92 | 0.50 | 7.00 | **20.32** | CER | Uncultured bacterium clone Ms-10-Fx11-2-091 | AB990033.1 | 99 | USA:Alaska - environmental_sample |
|  |  |  |  |  |  | CNR | Acetobacteraceae bacterium LX45 | KC921158.1 | 99 | Ginger foundation - soil |
| *Xenococcaceae* -6 | Cyanobacteria | 2.88 | **76.36** | 2.00 | - | CER | Uncultured cyanobacterium clone FQSS103 | EF522323.1 | 99 | Rocky Mountain - endolithic sandstone community |
|  |  |  |  |  |  | CNR | *Phormidium* sp. CCALA 726 | GQ504036.1 | 92 | Ny-Ålesund, Svalbard, Arctic |
| *Xenococcaceae* -2773 | Cyanobacteria | 2.54 | **159.23** | - | 0.25 | CER | Uncultured cyanobacterium clone FQSS103 | EF522323.1 | 98 | Rocky Mountain, USA - endolithic sandstone community |
|  |  |  |  |  |  | CNR | *Chroococcus* sp. VP2-07 | GQ504036.1 | 91 | Italy and Spain - fountains |
| *Sphingomonas* -15 | Proteobacteria | 2.15 | - | **147.50** | 132.00 | CER | Uncultured bacterium clone Bysf-47-Sf10-014 | KP296188.1 | 99 | Antarctic - surface seawater |
|  |  |  |  |  |  | CNR | *Sphingomonas* sp. UYEF32 | KU060875.1 | 99 | King George Island, Antarctica - exfoliation rock |
| *Sphingomonadaceae* -16 | Proteobacteria | 1.83 | 3.61 | **44.00** | 29.32 | CER | Uncultured bacterium clone Bysf-47-Sf10-014 | JX967335.1 | 99 | Norway - granite outcrop |
|  |  |  |  |  |  | CNR | *Blastomonas* sp. TW1 | AY704922.1 | 96 | Biofilm - drinking water |
| *Methylobacterium* -1508 | Proteobacteria | 1.38 | - | **184.50** | 9.00 | CER | Uncultured bacterium clone LIB079_B_C08 | KM852235.1 | 99 | Biofilm - drinking water |
|  |  |  |  |  |  | CNR | *Methylobacterium brachiatum* strain ZJ0902B96 | KU173699.1 | 98 | East China sea - surface seawaters |
| *Leptolyngbya* -290 | Cyanobacteria | 1.09 | **41.00** | - | - | CER | Uncultured cyanobacterium clone H-D14 | DQ181732.1 | 99 | East Antarctic lakes - microbial mat |
|  |  |  |  |  |  | CNR | *Phormidesmis* sp. LD30 5700 TP | LN849930.1 | 98 | Western Himalaya - biological soil crust |
| *Acetobacteraceae* -32 | Proteobacteria | 1.08 | - | **18.40** | 9.23 | CER | Uncultured bacterium QA4_30_153 | LC076735 | 98 | Cryoconite, Qaanaaq, Greenland |
|  |  |  |  |  |  | CNR | *Acetobacter pasteurianus* NBRC 3225 | AB680032.1 | 93 | Culture collection |
| *Leptolyngbya* -3 | Cyanobacteria | 0.75 | **92.80** | 44.00 | 45.35 | CER | Uncultured cyanobacterium clone LH16_269 | KM112118.1 | 99 | McMurdo Dry Valley Lakes, Antarctica - benthic mats |
|  |  |  |  |  |  | CNR | *Phormidesmis priestleyi* ANT.L66.1 | AY493581.1 | 99 | Antarctic lake - benthic microbial mat |
| *Leptolyngbya* -106 | Cyanobacteria | 0.51 | **141.64** | - | 6.67 | CER | Uncultured bacterium clone IT2-66 | KX247359.1 | 98 | Zhadang, China - glacier forefield |
|  |  |  |  |  |  | CNR | *Phormidesmis priestleyi* ANT.LG2.4 | AY493580.1 | 98 | Antarctic lake - benthic microbial mat |
| *Leptolyngbya* -76 | Cyanobacteria | 0.51 | **207.39** | - | 8.43 | CER | Cyanobacterium cWHL-1 | HQ230236.1 | 99 | Ward Hunt Island, Ward Hunt Lake, Canada - snow |
|  |  |  |  |  |  | CNR | *Phormidesmis priestleyi* ANT.LG2.4 | AY493580.1 | 98 | Antarctic lake - benthic microbial mat |
| *Methylobacterium* -6342 | Proteobacteria | 0.20 | - | 185.00 | **226.77** | CER | Uncultured alphaproteobacterium clone CN-2_B05 | EF219937.1 | 99 | Antarctica, unvegetated soil environments at Coal Nunatak |
|  |  |  |  |  |  | CNR | *Methylobacterium tardum* strain IHBB 11162 | KR085941.1 | 99 | Suraj Tal, Trans Himalayas - Lake Water |

## DISCUSSION

### Snow bacterial communities

Our results provide the first insights into the dynamics of bulk and potentially active communities of decaying snowpacks in the ablating zone of the ice sheet during the transition to bare ice. At the start of the study bacterial PSP is positive (Figure 2), however this rapidly declines for the remainder of the sampling period. In contrast, 16S rRNA gene copy numbers increase by 3-4 orders of magnitude for the remainder of the sampling period. This is likely due to the accumulation of biomass within decaying snow due to physical processes rather than biological growth (Björkman et al 2014). Melting snowpacks are physically and chemically dynamic environments, and it appears only a few lineages are able to maintain their populations in supraglacial snow as it decomposes to slush (Hell et al 2013), with other taxa being washed out. Indeed, 16S rRNA gene and 16S rRNA (cDNA) OTU profiles were highly significantly different over time (Supplementary Table 2).

Here, snowpack 16S rRNA gene copies greatly exceed 16S rRNA copy numbers, indicating the bulk community is likely to be exported as cells with low PSP. For example, the relative under-representation of Bacteroidetes in 16S rRNA (cDNA) raises the possibility that cellulose-degrading taxa become quiescent when dissociated from sources of complex organic carbon, for example supraglacial phototrophs (Smith et al 2016). It is therefore likely that the abundant groups of bacteria in decaying snow serve as sources of cellular carbon and nutrients rather than viable taxa capable of inoculating downstream habitats. The rare, high PSP *Methylobacterium* sp. OTUs detected represent an exception which will be discussed below.

Although most of the Greenland Ice Sheet is perennially covered with snow, few studies have examined the snowpack microbiology of the Greenland Ice Sheet (Cameron et al 2014). Moreover, the highly isolated setting of field sites coupled with the potential for contamination of low-biomass samples make such studies challenging. By establishing a field camp for the duration of the study, careful handling of samples and the sequencing of negative controls we were able to mitigate these limitations. Negative controls returned very small numbers of reads (Supplementary Table 3). Prominent groups of bacteria in our study were not represented in negative controls with the exception of seven reads matching *Phormidesmis priestleyi,* likely indicating post amplification carry-over of a dominant amplicon type at negligible levels compared to its abundance in field samples.

### Cryoconite bacterial communities

Lower copy numbers of 16S rRNA genes were amplified by qPCR compared to previous studies employing 16S rRNA gene qPCR based upon larger, wet-weight samples (Stibal et al 2015). However, the overall trends are consistent between both studies. Considering potential limitations in extraction efficiency and biases inherent in all PCR based analyses, we avoid treatment of qPCR data as absolute quantities of 16S rRNA genes or 16S rRNA in our samples and limit our comparison to trends within the dataset.

Exceptionally high ratios of 16S rRNA to rRNA genes were measured in cryoconite (Figure 2). Combined with amplicon sequencing data revealing cyanobacteria were overwhelmingly dominant within the 16S rRNA (cDNA) population (Figure 3, Figure 5) we interpret this as evidence of the high PSP of cyanobacteria within the cryoconite granules. Since filamentous cyanobacterial phototrophs such as *Phormidesmis priestleyi* are well known as ecosystem engineers (Cook et al 2015, Edwards et al 2014a) of cryoconite granules through their primary production and granule-building (Langford et al 2010), this is highly plausible.

Other work within the same field season at the same site lends support to our findings. Firstly, *Phormidesmis priestleyi* was isolated in culture and genome sequenced (Chrismas et al 2016) and secondly perturbation of cryoconite hole structure and microbial activity revealed *Phormidesmis* sp. employ sensitive photoadaptive mechanisms to optimize carbon sequestration in cryoconite holes (Cook et al 2016). Correspondingly, the prominence of OTUs extremely closely related to *Phormidesmis priestleyi* (Table 1), albeit assigned to *Leptolyngbya* (−3 and −76) within the high PSP (Figure 5) and keystone taxa (Table 1) of cryoconite granules extends insights from previous studies by linking specific *Phormidesmis* lineages with metabolic activities within Arctic cryoconite.

Importantly, previous analysis of 16S rRNA genes has resolved a single *Phormidesmis* OTU is cosmopolitan (Segawa et al 2017) within diverse Arctic glacial settings (Cook et al 2016, Edwards et al 2011, Gokul et al 2016, Uetake et al 2016). However, in our study, while one *Phormidesmis* OTU (*Leptolyngbya*-3) is likely to play a structural role, two other lineages (*Leptolyngbya*-76, *Leptolyngbya*-106) have PSP in gross excess to their biomass, indicated by contrasting trends in PSP and 16S rRNA gene relative abundance (Figure 5). The bulk bacterial community structure of cryoconite granules was stable over the course of the study, consistent with prior studies (Musilova et al 2015). However the *Phormidesmis* dominated 16S rRNA (cDNA) pool of the bacterial community of cryoconite changed over time, with highly significant changes revealed by PERMANOVA (Supplementary Table 2). Therefore, the potential for metabolic and structural niche differentiation among cryoconite *Phormidesmis* merits further investigation.

### Supraglacial stream water bacterial communities

Meltwater runoff from the Greenland Ice Sheet surface is thought to be a major contributor to sea level rise (Smith et al 2017). Although this meltwater is an important source of organic carbon and nutrients to downstream ecosystems (Bhatia et al 2013a, Bhatia et al 2013b, Hawkings et al 2016, Musilova et al 2017) and the microbial fluxes in outflows of the Greenland Ice Sheet have been studied (Cameron et al 2017, Dubnick et al 2017), the absence of data on microbial export from the Greenland Ice Sheet surface represents a critical lacuna in our understanding of the Greenland Ice Sheet ecosystem. Within this study, this is addressed by 16S rRNA gene and 16S rRNA (cDNA) qPCR and sequence data which reveal the export of microbial groups prevalent in snow and cryoconite in meltwater from three ephemeral supraglacial streams.

Quantitative PCR reveals the stream water bacterial community in the first week of the study contains approximately equal copy numbers of bacterial 16S rRNA genes and 16S rRNA (cDNA) resulting in a bacterial PSP of 0.97. By the second week, both genes and rRNA have increased their average copy number by two orders of magnitude. Highly significant differences were observed in the community structure of bulk, but not active stream bacterial communities over time (Supplementary Table 2). Only one profile each of the bulk and active communities could be analysed from week one, but both were strongly dominated by Alphaproteobacteria. Subsequent weeks are marked by a more diverse bulk bacterial community, although *Alphaproteobacteria* were highly dominant in the potentially active bacterial community throughout. Sphingomonadaceae, Acetobacteraceae, and Rickettsiaceae are prevalent in the high PSP *Alphaproteobacteria* detected in stream water, but the highest PSP taxon is *Methylobacterium*-1, which is also a high-betweenness putative keystone species.

The presence of cyanobacterial taxa associated with cryoconite, including *Leptolyngbya-3* and *Leptolyngbya-*76 in stream water indicates the export of potentially active primary producers. It is likely these cyanobacteria originate from biomass sheared from cryoconite granules either present in cryoconite holes, as distributed cryoconite on the ice surface or in stream cryoconite (so-called “hydroconite”; (Hodson et al 2007). It is likely these phototrophs represent sources of highly bioavailable dissolved organic carbon exported from the glacier surface (Musilova et al 2017).

### High Protein Synthesis Potential Rare Taxa in the Dark Zone of Greenland

A consistent pattern for all three habitats sampled was that bulk and potentially active communities of snow, cryoconite and stream habitats were highly significantly different (Supplementary Table 2, Figure 3). Furthermore, a small subset of taxa were consistently over-represented in 16S rRNA (cDNA) compared to their corresponding 16S rRNA gene relative abundances, most notably the Cyanobacteria in cryoconite and Alphaproteobacteria in snow and stream habitats. Other taxa were typically under-represented. The majority of taxa present within the surface of the Dark Zone appear to exhibit low protein synthesis potential. This may be due to resource limitation, dormancy or the detection of DNA associated with dead cells (Blazewicz et al 2013). Each of the above scenarios has important ecological implications. Firstly in terms of maintaining a pool of organisms which may respond to stimuli such as allochthonous resources, and secondly, the maintenance of a “seedbank” of conditionally viable cells (Lennon and Jones 2011), or at the very least, the contribution of carbon and nutrients in otherwise oligotrophic environments in the form of necromass. Differentiation between these scenarios is beyond the scope of 16S rRNA gene analyses (Blazewicz et al 2013) and the potential for “active” taxa to be mis-classified as “dormant” by 16S rRNA (cDNA) / 16S rRNA gene comparison has been described in computational simulations (Steven et al 2017).

Nevertheless, a further notable trend which was consistent across all three habitats was a marked prominence of rare taxa among those with high PSP. Such patterns have been observed in other, comparable contexts, for example within proglacial streams in the European Alps (Wilhelm et al 2014). In those systems, such trends have been considered hallmarks of habitats exhibiting severe fluctuations in their environmental conditions such as temperature, discharge or solar radiation at timescales briefer than the doubling time of the resident community (Lennon and Jones 2011, Wilhelm et al 2014). Here, no overall trends were apparent in terms of the influence of meteorological parameters from week to week, however when monitored continuously (Figure 1) profound oscillations in temperature, incoming short- and long-wave radiation, air temperature, energy flux and melt intensity are apparent at diurnal to sub-weekly periodicity. Considering the sluggish growth of organisms at temperatures at or near freezing (Anesio et al 2010), it is likely that such oscillations create rapidly changing niche spaces at timescales shorter than community doubling times. It is therefore likely that organisms exhibit high PSP relative to their biomass when they are able to maintain activity but not achieve net population growth in the face of unstable fitness.

Consequently, through their metabolic activities, high PSP rare taxa may be disproportionately influential within their communities, fitting the definition of keystone taxa. This is coherent with the corresponding prevalence of high PSP rare taxa among putative keystone taxa identified by betweenness (Table 1). Gokul et al (Gokul et al) previously identified supraglacial keystone taxa in the cryoconite communities of a High Arctic ice cap, and the present study extends the case that specific rare taxa exert disproportionate influence on the bacterial communities of glacier surfaces through maintaining high levels of metabolic potential. Table 1 reports these taxa typically possess very close relatives (either as environmental sequences or named isolates) in a diverse range of habitats within the global cryosphere, with two implications. Firstly, this lends pragmatic support to their likely authenticity within the communities of the Greenland Ice Sheet Dark Zone, but secondly, the inference is that adaptations resulting in disproportionately high PSP may be common among cosmopolitan species in the polar and alpine regions.

### Implications for biogeochemical cycling

The prominence of OTUs assigned to *Methylobacterium* in the high PSP taxa of snow and stream water communities is very apparent. In particular, the exceptionally high PSP shown by the *Methylobacterium-*1 OTU is striking for both habitat types, with other related OTUs (*Methylobacterium-*6342 and *Methylobacterium*-1508) showing very high PSP. *Methylobacterium*-1 is well represented within the snowpack 16S rRNA gene profiles of week 1, but then shows disproportionately high PSP in following weeks before its loss from the snowpack community by the fifth week. This would suggest continued protein synthesis potential is maintained for some time in spite of its rapidly diminished population size within the snowpack.

Combined, this pattern of high PSP taxa affiliated to the genus *Methylobacterium* merits further consideration. *Methylobacterium* are well known facultative methylotrophs (Chistoserdova et al 2003, Chistoserdova 2011), raising the prospect of methyl metabolism as a hitherto unappreciated metabolic strategy on the Greenland Ice Sheet. Redeker et al (Redeker et al 2017) provided direct evidence of trace gas metabolism in polar snowpacks through the cycling of methyl halides and dimethyl sulphides. Although Redeker et al (2017) did not explore the diversity of microbes associated with methyl cycling, it is possible that *Methylobacterium* in the decaying snowpacks of the Dark Zone are involved in cycling of climate-relevant trace gases.

The relevance of disproportionately high PSP *Methylobacterium* to biogeochemical cycling in the Dark Zone is further amplified when their status as the dominant 16S rRNA (cDNA) taxon in meltwater exports is considered. Supraglacial meltwater from the Dark Zone is typically routed via surface streams into moulins to the bed of the Greenland Ice Sheet, an environment conducive for methane cycling(Yang and Smith 2016). In catchments fed by meltwater from the Dark Zone, variable rates of methanogenesis and methane oxidation have been observed, with potential impacts for the global methane cycle (Dieser et al 2014, Lamarche-Gagnon et al 2019). The discharge of highly oxygenated supraglacial meltwater into the subglacial environment is strongly associated with the cessation of methanogenesis and consumption of methane via aerobic oxidation (Dieser et al 2014). The export of high PSP *Methylobacterium* from the Dark Zone surface in this meltwater, as detected here, raises the prospect that surface-derived taxa inoculating the bed of the Greenland Ice Sheet play a role within a subglacial consortium nourished by the oxidation of methane. Further investigations focused on the fate of supraglacial microbiota transferred to the subglacial ecosystem could reveal whether this process is sufficient to mitigate the subglacial synthesis of this potent greenhouse gas.

## Conclusions

In summary, 16S rRNA gene and rRNA (cDNA) quantification and sequencing of snow, cryoconite and stream water bacterial communities from the Dark Zone of the Greenland Ice Sheet was conducted at weekly intervals during the melt season of 2014. Recently, attention has been focused on the albedo-reducing properties of microbial consortia within the Dark Zone (Williamson et al 2019) highlighting the importance of microbial interactions in the future of the Greenland Ice Sheet. Our study addresses the related question of microbial community dynamics, and reveals that rare taxa appear to be disproportionately active. Notably, these taxa appear central to the structure of their communities and may play under-appreciated roles within the carbon cycle of the Greenland Ice Sheet. The presence of high-PSP rare taxa within *Methylobacterium* in melting snow and stream-water raises the prospect of supraglacial methyl compound cycling and export to the subglacial ecosystem. Our study represents a targeted locus amplicon sequencing approach, which in future could be complemented with genome-resolved metagenomics and direct process measurements of carbon cycling and export in both Dark Zone surface and connected downstream habitats. This would further elucidate the connections between these communities, climate change and impacts on downstream riverine and marine ecosystems from the most expansive supraglacial bare ice habitat in the Northern Hemisphere.

## ACKNOWLEDGEMENTS

This manuscript is dedicated to the memory of Kathi Hell (1985-2019) whose earlier work with us on dynamic Arctic snow microbiomes helped inspire the study described herein. Fieldwork and laboratory analyses were supported by Royal Society grant RG130314 to AE and TI-F while JKG was supported by a South African National Research Foundation Fellowship. Sequencing was performed using BBSRC funded facilities for the analyses described herein. KAC acknowledges funding from the European Union’s Horizon 2020 research and innovation programme under the Marie Skłodowska-Curie grant agreement No 663830. Financial support was also provided to KAC by the Welsh Government and Higher Education Funding Council for Wales through the Sêr Cymru National Research Network for Low Carbon, Energy and Environment. AE is grateful for Leverhulme Trust Research Fellowship RF-2017-652\2 and NERC NE/S1001034/1 which eased completion of the work.

## The authors declare no conflict of interest

## Supplementary Methods

### Sample archiving

Other researchers have documented the half-life of the cellular RNA pool (which is heavily dominated by rRNA e.g. Moran *et al*. 2012) of glacial bacteria at the low temperatures typical of ice surfaces is less than one day (Segawa *et al*. 2014), with the implication that rRNA turnover rates are sufficient at the temporal resolution of this study to capture meaningful changes in protein synthesis potential well within the expected community doubling time (Anesio *et al*. 2010). However, to prevent sample degradation, this necessitates careful sample archiving to stabilize the rRNA and genomic DNA pools of the collected biomass. Therefore, due to the remote location of the field camp, in line with established procedures (e.g. Stibal *et al*. 2015, Cameron *et al*. 2015), Soil Lifeguard RNA/DNA preservative (MO BIO Laboratories) stabilized samples were stored dark, within crushed ice, prior to transfer to −20°C storage at the Kangerlussuaq International Science Support field laboratory, within three weeks of collection, followed by −80°C archiving at the Aberystwyth laboratory until further processing within four months.

Specifically, in the field, cryoconite samples were aspirated into sterile microcentrifuge tubes and immersed immediately in Soil Lifeguard RNA/DNA preservative (MO BIO Laboratories). Snow samples representing 1.2-1.5L water equivalent were collected in sterile whirlpaks (Nasco, Inc.) and melted in the dark at +10°C while meltwater from supraglacial stream meltwater (ca. 2L) was collected in whirlpaks. Melted snow was filtered through 0.22 µm Sterivex GP polyethersulfone filters (Millipore) and chemically preserved within 12 hours of sampling, and stream water was filtered and chemically preserved within 6 hours of sampling. The Sterivex filters were connected to tubing which was thoroughly rinsed using 10% HCl and 70% ethanol to remove potential contaminants from the tubing entering the whirpak bag. Sterivex filters were filled with 2 mL Soil Lifeguard RNA/DNA preservative (MO BIO Laboratories).

### Nucleic Acid Extraction

All pre-PCR procedures were conducted aseptically in a bleach-disinfected laminar flow hood with certified DNA/RNA free plasticware including aerosol resistant tips. Negative extraction controls were prepared using sterivex cartridges and blank cryoconite extraction.

Total nucleic acids were co-extracted from cryoconite using the PowerBiofilm^™^ RNA Isolation Kit (MO BIO Laboratories) according to the manufacturer’s instructions. 0.2g ±0.05g of wet sediment was used in each extraction. Nucleic acids from snow and stream samples were co-extracted using the PowerWater® Sterivex™ RNA Kit (MO BIO Laboratories) according to the manufacturer’s protocol with minor adjustments for DNA extraction recommended by MO BIO Laboratories (Dr Emelia DeForce, personal communication). Specifically, these steps required the addition of 20 µL beta-mercaptoethanol for every 880 µL of solution ST1B, the additional incubation of the sterivex filter at 70°C for 10 minutes during lysis and the addition of an equal volume of 100% ethanol to buffer ST4. All RNA samples were subjected to DNase treatment using the RTS DNase^™^ Kit (MO BIO Laboratories) according to the manufacturer’s instructions. Aliquots of each RNA sample were then used as template in first strand cDNA synthesis via reverse transcription with SuperScript™ III Reverse Transcriptase (Invitrogen) and universal 16S rRNA gene region primer 1389R, according to the manufacturer’s instructions, for use in downstream PCR applications.

### 16S ribosomal RNA (cDNA) and 16S ribosomal RNA gene quantitative PCR

To estimate 16S rRNA gene and 16S rRNA abundance within the samples, two microliters of each DNA extract or reverse transcriptase product was used as template in 20 µL reactions consisting of 1× SensiFast SYBR (no ROX) qPCR mixture (BioLine, Ltd) and 16S rRNA primers 27F (5’-AGAGTTTGATCCTGGCTCAG) and 1389R (5’-ACGGGCGGTGTGTACAAG) amplified for 35 cycles in a Mic real time PCR cycler (BioMolecularSystems, Inc.) The copy number of gene and 16S rRNA molecules was estimated from a plasmid-borne 27F-1389R standard, and converted to gram dry weight cryoconite or meltwater stream volume or water equivalent volume of snow (by reference to the volume of samples filtered) respectively. We note the uncertainties associated with extraction efficiencies for different microbial taxa within environmental samples subjected to nucleic acid extraction, therefore we present the copy number of 16S rRNA genes and 16S rRNA as the *amplifiable* copy number per unit sample.

### 16S ribosomal RNA gene amplicon sequencing

Cryoconite, snow and stream DNA and RNA samples were prepared for paired end MiSeq sequencing (Illumina). Using a 3 stage PCR method, the bacterial 16S rRNA V3-V4 hypervariable region was amplified for the attachment of Nextera XT dual indices and Illumina sequencing adapters.

The primers used for 16S rRNA gene specific amplification were universal 16S primers 27F and 1389R using cycling conditions of 95 °C for 5 mins (initial denaturation); 30 cycles: 95 °C for 30 secs (denaturation), 60 °C for 30 secs (annealing), 72 °C for 45 secs (extension); 72 °C for 7 mins (final extension). Amplicons produced were used as template for nested PCRs with Nextera primers (5’-TCGTCGGCAGCGTCAGATGTGTATAAGAGACAG-3’ and 5’- GTCTCGTGGGCTCGGAGATGTGTATAAGAGACAG-3’) (Klindworth et al, 2013) in 12.5µL Platinum Taq Mix PCR reactions, as per manufacturer’s instructions, using cycling conditions of 95°C for 3 mins (initial denaturation); 25 cycles: 95°C for 30s (denaturation), 55°C for 30s (annealing), 72°C for 30s (extension); 72°C for 5 mins (final extension). Amplification was confirmed by gel electrophoresis on 1% agarose before products were used as template in the third PCR using Platinum Taq Mix and the Nextera XT Index Kit (Illumina) to obtain unique combinations of the N7 and S5 primer barcodes per sample (SUPPLEMENTARY TABLE 1). Cycling conditions were 95° for 3 mins (initial denaturation); 8 cycles of 95°C for 30s (denaturation), 55°C for 30s (annealing), 72°C for 30s (extension); 72°C for 5 mins (final extension). Amplicons were purified using the Montage™ PCR plate filtration system (Millipore) prior to quantification on an Epoch™ spectrophotometer (BioTek). Products were then normalised to the lowest common concentration for each library (DNA 5ng/µL; cDNA 0.8ng/µL) and pooled before electrophoresis on 1% agarose gel for purification via gel extraction using the Qiagen Gel Extraction Kit (Qiagen) as per manufacturer’s instructions. Libraries were sequenced at the IBERS Translational Sequencing Facility (IBERS Phenomics Centre, Gogerddan, Aberystwyth, Ceredigion SY23 3EE) on the Illumina MiSeq® platform (Illumina Inc., San Diego, CA, USA) using the MiSeq® Reagent Kit, version 3 (Illumina Inc., San Diego, CA, USA). Initial processing of Illumina MiSeq® sequence data was performed in BaseSpace® (Illumina Inc., San Diego, CA, USA). Data were demultiplexed and the indices removed. Reads were trimmed to remove read-through into the adaptor sequence at the 3’ end. Sequence files were generated in the fastq format and imported into QIIME 1.9.0 for merging of paired end reads.

### Sequence Processing and Analysis

Resulting sequences were quality filtered and processed in QIIME 1.9.0 (Caporaso *et al*., 2010) using default quality filters unless otherwise stated. Paired end sequences were joined and probed for chimeras using USearch6.1 (Edgar, 2010) which were then removed. Operational taxonomic units (OTUs) were then assigned at 97% identity using a reference based UCLUST algorithm (Edgar, 2010) and the Greengenes 13_8 database, August 2013 version (DeSantis *et al*., 2006) before OTU tables were generated, and processed for rarefaction curves and alpha diversity indices. Multivariate analysis was performed in PRIMER 6/PERMANOVA+ (PRIMER-E Ltd, Plymouth UK). Permutational multivariate analysis of variation (PERMANOVA), principal coordinates analysis (PCO) were performed as described previously (Edwards *et al*., 2014) with Bray Curtis transformations at the fourth root used for OTU relative abundance. One-way ANOVA was performed in Minitab 17. For samples with both 16S rRNA gene and 16S rRNA profiles, protein synthesis potential (sensu Denef *et al*., 2016) was defined as the ratio of the relative abundances of each OTU. Community analysis using pairwise Spearman correlations in R was able to identify OTUs with high *betweenness* centrality (shortest number of paths between any two other OTUs passing through that OTU) (Peura et al, 2015), high *degree* centrality (the number of associations an OTU has) and high *closeness* centrality (the average distance between two OTUs) measures (Berry and Widder, 2014). These taxa play a central role in the ecosystem and are recognised as “bottleneck or keystone taxa” (Paine, 1966, Mills et al, 1993, Cottee-Jone and Whittaker, 2012). Sequences are available on EBI-SRA under the study accession number PRJNA318626.

---

**SUPPLEMENTARY TABLE 1:**
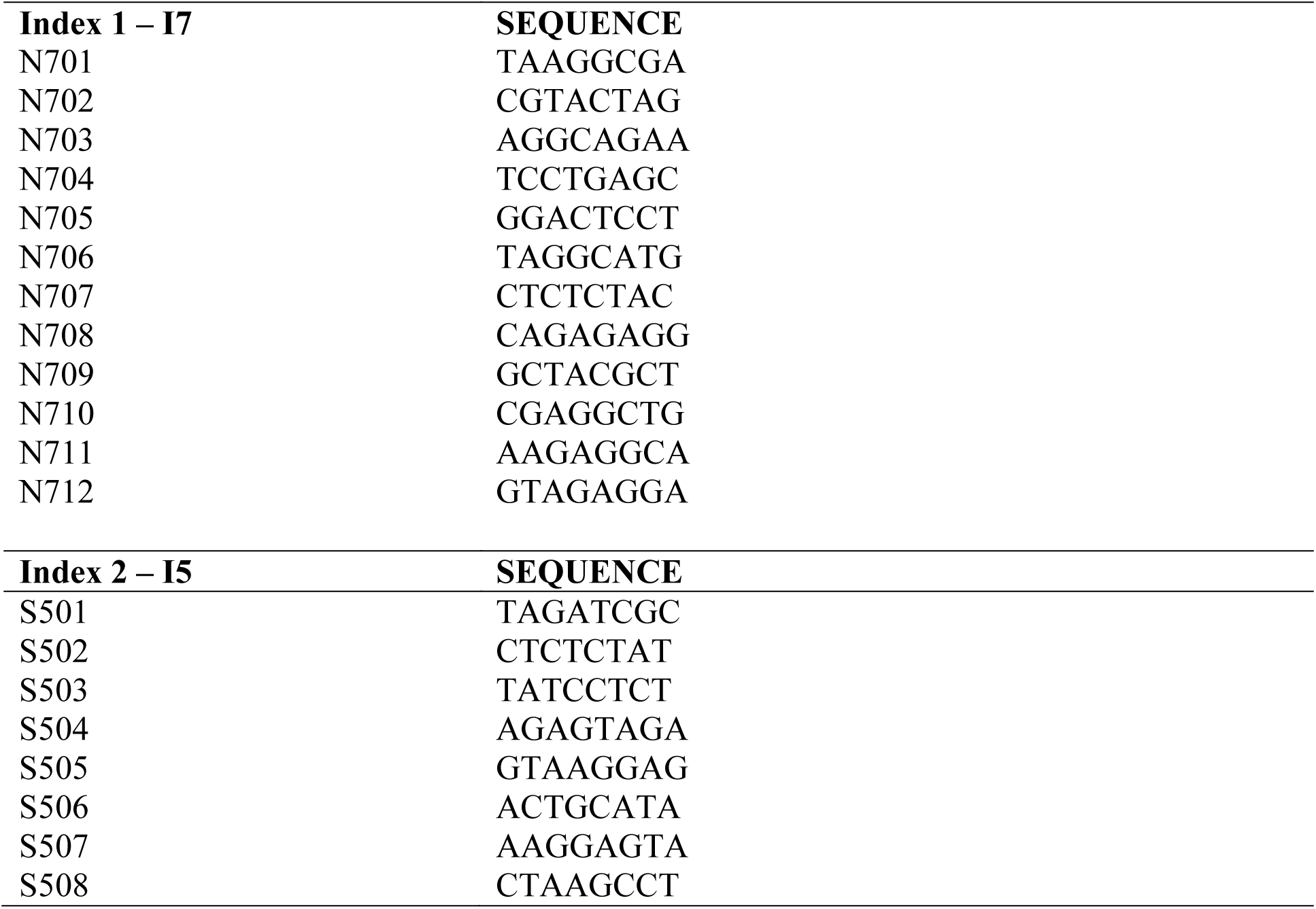
Dual Index Primers for the 96 sample Nextera XT Index Kit.

**SUPPLEMENTARY TABLE 2:** Permutational analysis of variance (PERMANOVA) across 16S rRNA gene and 16S rRNA (“nucleic acid”), day of year ad cryoconite, snow,and stream habitats.

| Test | Pseudo $F$ | $p$ |
| --- | --- | --- |
| Habitats | 13.796 | 0.0001 |
| Cryoconite DNA vs RNA | 22.536 | 0.0001 |
| Cryoconite DNA over time | 1.0092 | 0.4477 |
| Cryoconite RNA over time | 1.6963 | 0.0254 |
| Snow DNA vs RNA | 5.3008 | 0.0001 |
| Snow DNA over time | 2.6525 | 0.0001 |
| Snow RNA over time | 2.9937 | 0.0003 |
| Stream DNA vs RNA | 7.1009 | 0.0001 |
| Stream DNA over time | 2.4599 | 0.0041 |
| Stream RNA over time | 1.4237 | 0.1069 |

**SUPPLEMENTARY TABLE 3:** Identity and abundance of contaminating sequences detected in DNA extraction controls.

| OTU | Greengenes ID | No of reads in NTC | BLAST ID | Accession No | %ID |
| --- | --- | --- | --- | --- | --- |
| denovo-87257 | *g_Salinibacterium* | 16 | *Leifsonia* sp. URHE0078 | LN876553.1 | 99 |
| denovo-114112 | *f_Microbacteriaceae* | 12 | *Salinibacterium amurskyense* IC A9 | KU925169.1 | 98 |
| denovo-125684 | *o_Streptophyta* | 11 | *Roya obtusa* | KU646496.1 | 97 |
| denovo-90892 | *g_Leptolyngbya* | 7 | *Phormidesmis priestleyi* ANT.L66.1 | AY493581.1 | 99 |
| denovo-96273 | *g_Kocuria* | 4 | *Kocuria subflava* | NR_144586.1 | 99 |
| denovo-67624 | *f_Sphingomonadaceae* | 4 | *Polymorphobacter* sp. Ap23E | KX990242.1 | 99 |
| denovo-106624 | *f_Microbacteriaceae* | 3 | *Salinibacterium amurskyense* IC A9 | KU925169.1 | 98 |

**SUPPLEMENTARY TABLE 4:**
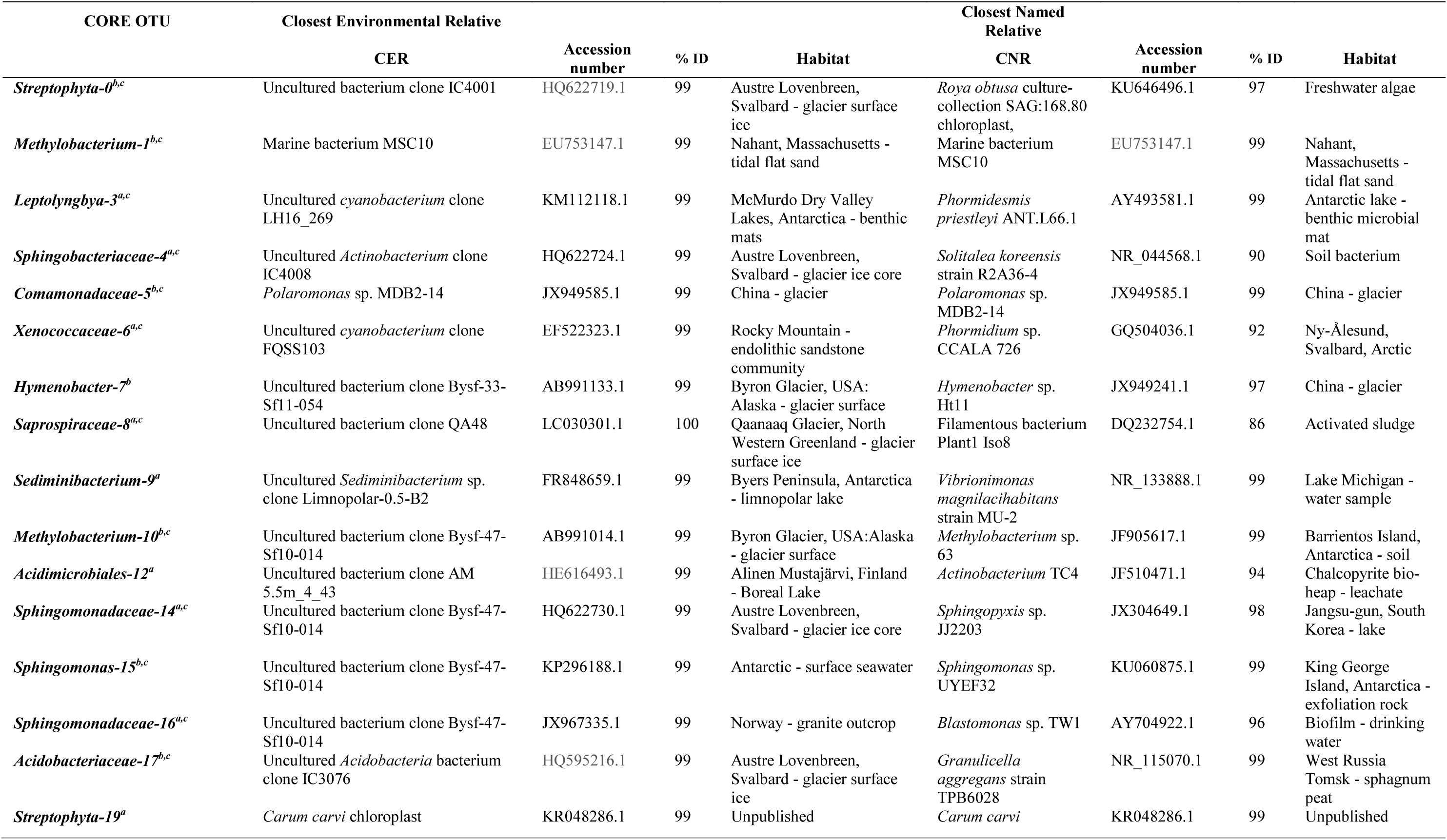
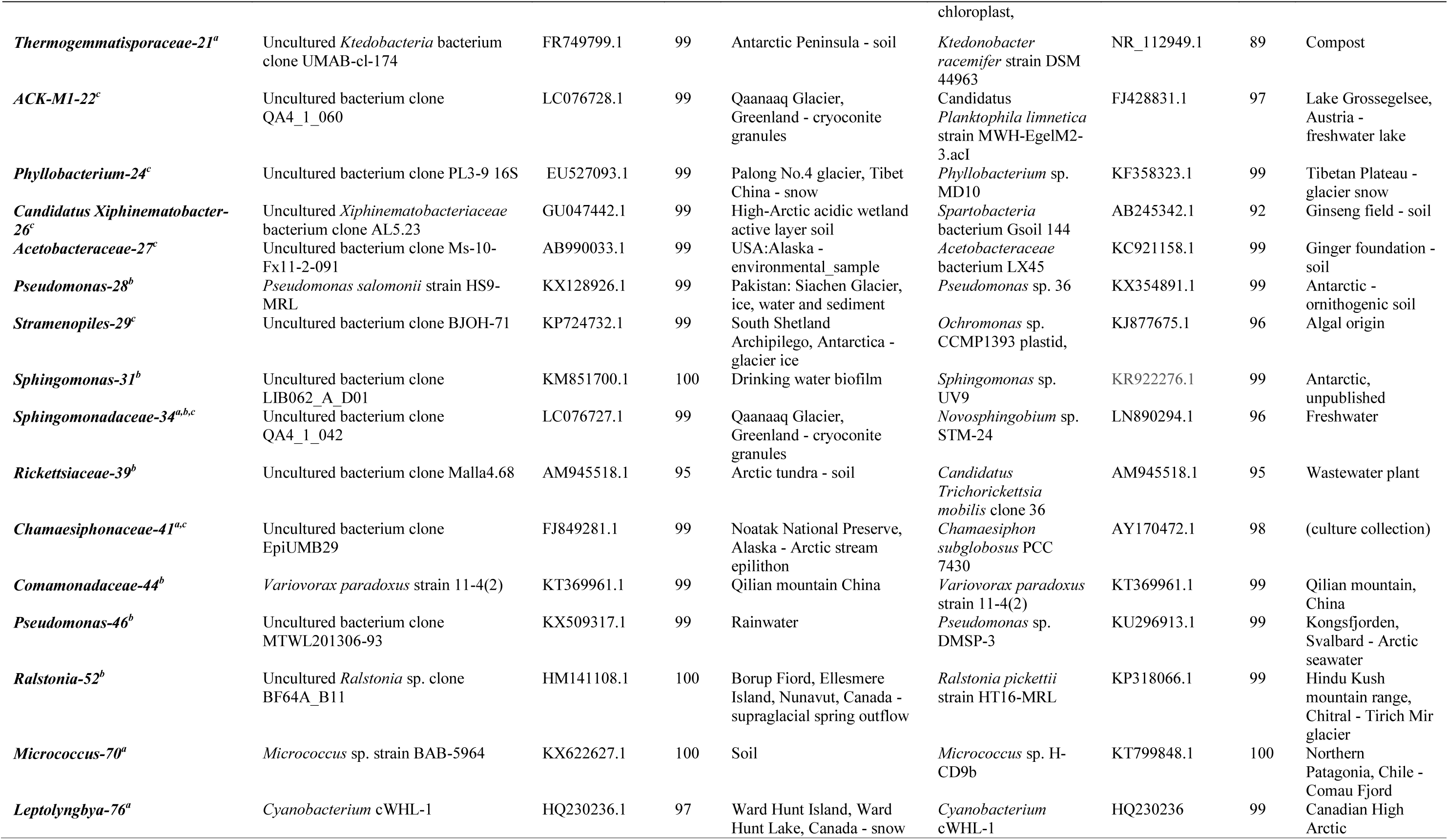
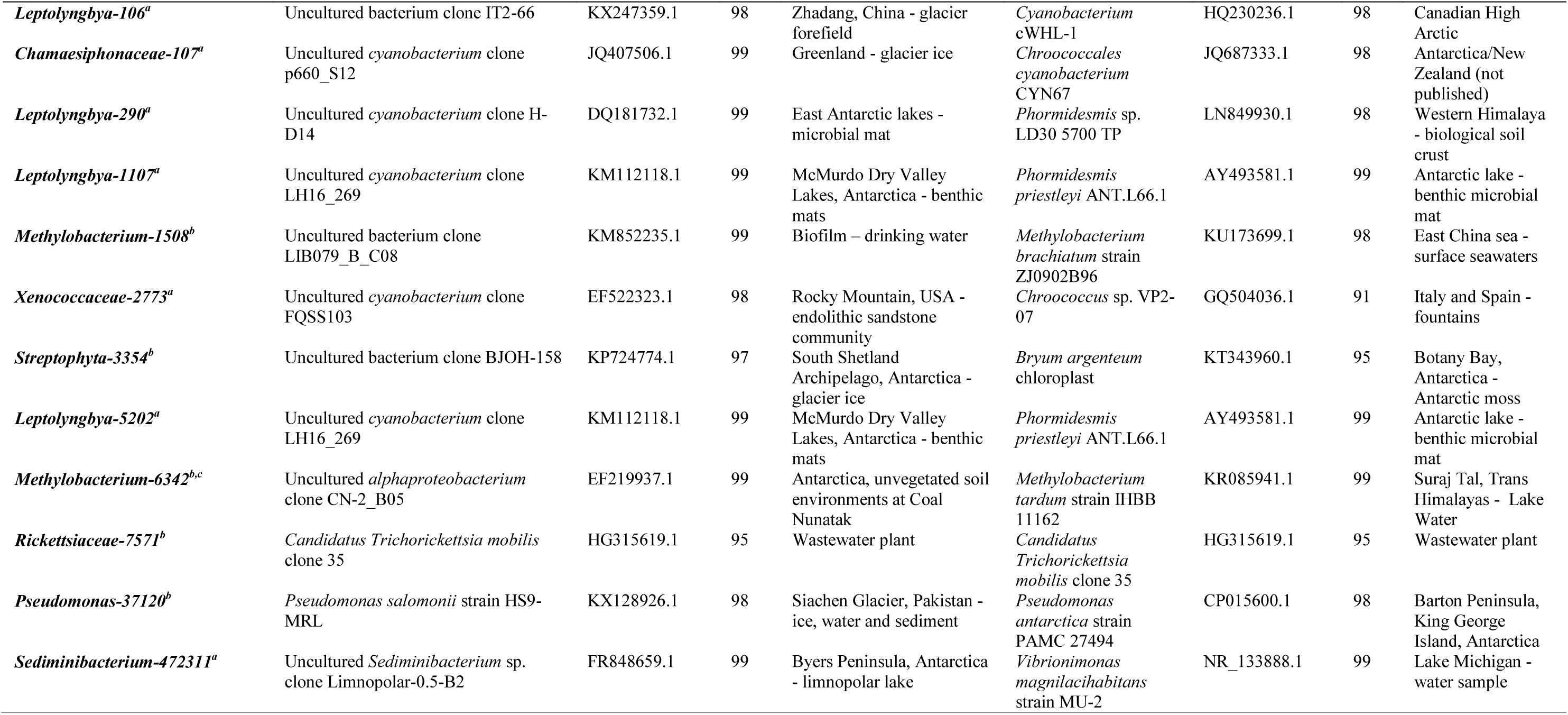
The identity of top OTUs present in cryoconite^a^, snow^b^ and stream^c^ water contributing to 99% of relative abundance per habitat.

